# Boundary stacking interactions enable cross-TAD enhancer-promoter communication during limb development

**DOI:** 10.1101/2023.02.06.527380

**Authors:** Tzu-Chiao Hung, David M. Kingsley, Alistair Boettiger

## Abstract

While long-range enhancers and their target promoters are frequently contained within a TAD, many developmentally important genes have their promoter and enhancers within different TADs. Hypotheses about molecular mechanisms enabling such cross-TAD interactions remain to be assessed. To test these hypotheses, we use Optical Reconstruction of Chromatin Architecture (ORCA) to characterize the conformations of the *Pitx1* locus on thousands of single chromosomes in developing mouse limbs. Our data supports a model in which neighboring boundaries are stacked with each other as a result of loop-extrusion, bringing boundary-proximal *cis*-elements into contact. This stacking interaction also explains the appearance of architectural stripes in the population average maps (e.g. Hi-C data). Through molecular dynamics simulations, we further propose that increasing boundary strengths facilitates the formation of the stacked boundary conformation, counter-intuitively facilitating border bypass. This work provides a revised view of the TAD borders’ function, both facilitating as well as preventing *cis*-regulatory interactions, and introduces a framework to distinguish border-crossing from border-respecting enhancer-promoter pairs.

## Introduction

The mammalian genome is partitioned in its 3D organization into topologically associating domains (TADs), regions in which intra-domain interactions are substantially more common than interdomain interactions ^1–4^. Known long range *cis*-regulatory interactions, most notably between enhancers and their target promoters, frequently lie within the same TAD, and in a multitude of cases mutations which disrupt this TAD organization result in corresponding disruptions to normal enhancer promoter contact and changes in gene expression ^5–9^.

Nonetheless, several developmentally important enhancers are separated by TAD borders and CTCF bound insulators from the genes they control. These include genes with key roles in axial patterning ^10^, hematopoiesis ^11^ and limb patterning ^12,13^. As candidate enhancer-promoter (E-P) interactions are frequently hypothesized and tested based on linear proximity to a gene, guided by previously mapped TAD boundaries, and studied in cell culture, it is likely that the genome-wide frequency of TAD border-bypassing E-P pairs is underestimated. The known examples, just mentioned, are among some of the longer-range acting enhancers and have preferentially been identified by low-throughput phenotype-driven studies in developmental contexts where cell culture models are often lacking. Being able to distinguish these border-crossing from border-respecting enhancers is an essential feature of understanding *cis*-regulation. A notable case of this border bypass occurs in the regulation of the gene *Pitx1*, where one of its major enhancers, *Pen*, is located no less than three TADs away (**Fig. 1a, b**).

**Fig. 1.**
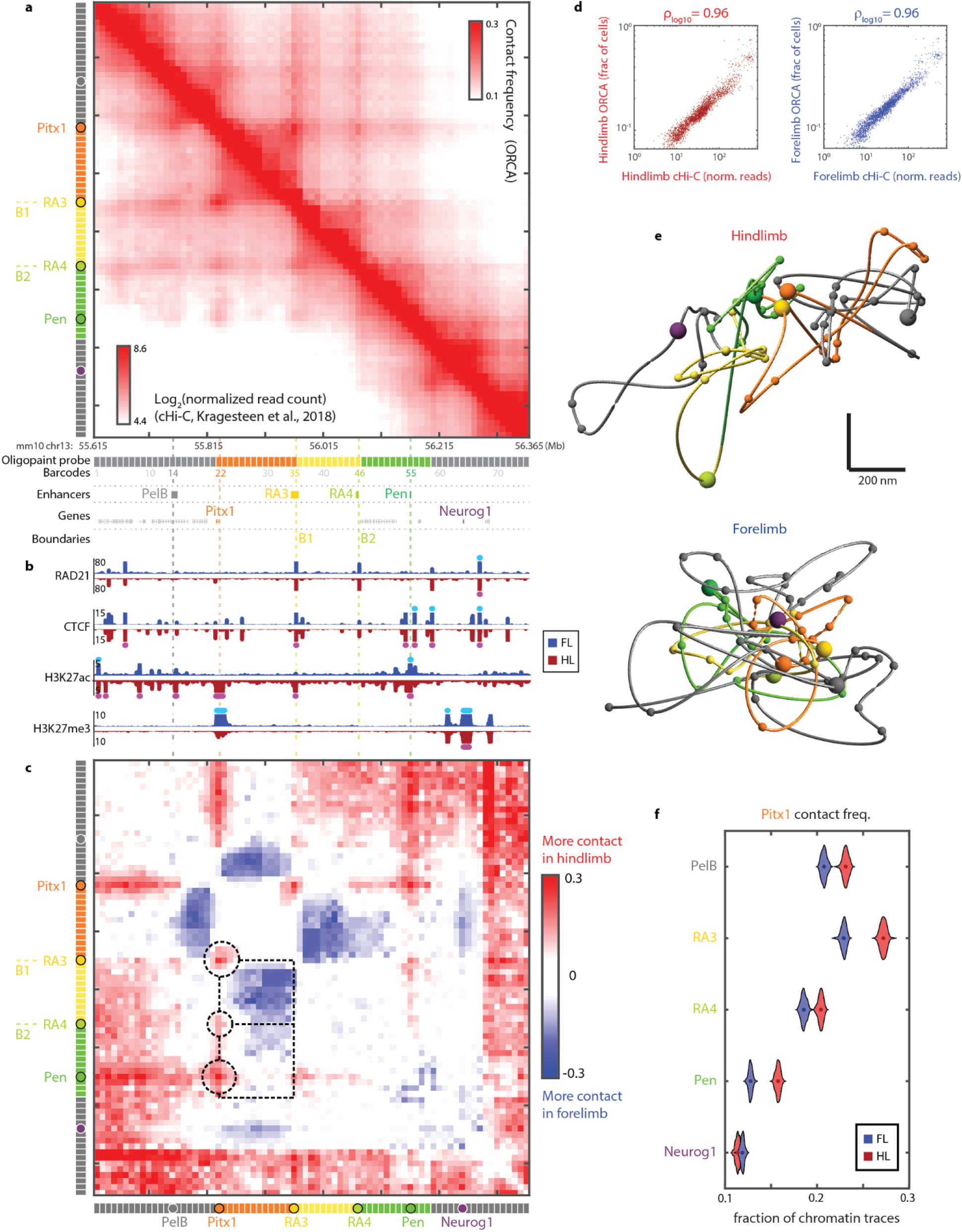
Chromatin conformation at *Pitx1* domain examined by ORCA agrees with published cHi-C data. **a,** Chromatin conformation at *Pitx1* domain in hindlimb cells as detected by ORCA (upper-right half, from 44,966 individual chromatin traces) and cHi-C ^18^ (lower-left half). **b,** ChIP-seq data from E10.5 limb buds (adapted from ^21^). Blue: Forelimb data; Red: Hindlimb data. The tracks are thickened to increase visibility. Signals higher than the y-axis limit are marked with bright turquoise and magenta for FL and HL, respectively. **c,** Difference in contact frequency, shown as the log2 ratio of hindlimb to forelimb. See text for discussion on the paradox highlighted by the dashed boxes and circles. **d,** Correlation of cHi-C and ORCA from hindlimb and forelimb. **e,** Representative reconstructed chromatin traces of the *Pitx1* chromosomal domain from hindlimb and forelimb visualized using ORCA. Spheres represent important genes and enhancers. See colorbar in **a** or **c** for reference. **f,** Violin plots quantifying the difference in contact between Pitx1 promoter and the indicated elements. All comparisons are statistically significant. (p = 2.6e-34, p = 2.6e-34, p = 2.8e-13, p = 2.6e-34, p = 2.6e-34, Wilcoxon test.)

Several properties make *Pitx1* ideal for studying border bypass. Whereas enhancer redundancy/shadow enhancers can often mask the regulatory contributions of individual enhancers and functional linking between enhancers and promoters ^14,15^, the *Pitx1* distal enhancer, called *Pen*, has obvious effects on both gene expression and morphological phenotypes. Indeed, studies in human patients first implicated the region around *Pen* in regulation of limb specific expression of *Pitx1* ^16,17^. Subsequent work with mouse genetic models found deletion of *Pen* caused a 45% reduction in *Pitx1* hindlimb expression, and increased incidence of clubfoot, consistent with reduced *Pitx1* function ^18^. In addition to the multi-TAD crossing *Pen*, two other hind-limb specific enhancers of *Pitx1* have also been characterized, which provide intriguing comparison cases. 5’ of *Pitx1* lies the RA3/PDE enhancer, which crosses no TAD borders in interacting with *Pitx1* ^19^. Meanwhile 3’ of *Pitx1*, in the neighboring TAD, lies the *PelB* enhancer in the intron of a neighboring unrelated gene ^20^. This enhancer is conserved through fish, but must navigate one border to interact with *Pitx1*.

Another attractive feature of *Pitx1-Pen* border bypass is its tissue-specificity. Though *Pitx1* is primarily expressed in hindlimb and not forelimb, the isolated *Pen* enhancer drives expression in both tissues when cloned and tested in a small transgenic reporter construct ^18^. Furthermore, the *Pen* enhancer in the native *Pitx1* locus carries the H3K27ac mark associated with active enhancers in both hindlimbs and forelimbs (**Fig. 1b ^21^)**. However, recent capture-Hi-C (cHi-C) experiments from mouse limb revealed quantifiably higher interaction frequency between *Pen* and *Pitx1* in the hindlimb compared to forelimb tissues ^18^. Notably, this work also showed that this difference was largely reduced when genetic aberrations removed a TAD boundary between *Pen* and *Pitx1* and brought them closer in linear distance, either through deletion or inversion, which led to increased *Pen-Pitx1* interaction, ectopic *Pitx1* expression in the developing forelimb, and corresponding forelimb to hindlimb developmental transformations ^18^. Thus, the hindlimb-specific interaction topology appears to account for the normal cell-type specific gene expression, but how this contact is achieved across multiple TAD boundaries remains a mystery. To understand the requirements for border bypass, here we compare how the same *Pitx1* chromosomal domain differs in its 3D conformations between forelimb and hindlimb, where border bypass is prohibited in the former but enabled in the latter.

We employ Optical Reconstruction of Chromatin Architecture ^22,23^ to visualize the 3D organization of the *Pitx1* chromosomal domain at single-chromatin resolution in developing limbs. This microscopy based approach, similar to other recent studies ^24–28^, allows the 3D chromatin trajectory to be visualized from individual cells, and thus has the potential to reveal what folding behavior enables hindlimb cells to achieve cross-TAD border enhancer-promoter interaction. By comparing 3D conformation of the *Pitx1* domain in the otherwise similar tissues of hindlimb and forelimb, we aimed to uncover the molecular mechanisms which enable functional bypass of TAD borders.

## Results

To visualize the *Pitx1* chromosomal domain, we designed Oligopaint probes tiling a 750 kb region around *Pitx1*, encompassing the downstream *PelB* enhancer, upstream enhancers RA3/PDE and *Pen*, and an unrelated gene *Neurog1* in the distal polycomb-silenced region (**Fig. 1a**). These probes span the 2 CTCF marked TAD borders, which we term B1 and B2, respectively, between *Pitx1* and *Pen* (**Fig. 1a, b**). We hybridized these probes to developing forelimbs and hindlimbs from E11.5 - E12.5 C57BL/6J mouse embryos, and visualized nearly 100,000 chromosome traces at 10 kb resolution, with each trace containing 75 3D coordinates, across 5 individual experiments (**Supplementary Table 1**).

The pairwise contact frequency (where two positions were found in proximity i.e., within 200 nm of one another) agreed closely with the corresponding frequency at which ligated reads were detected in previous work using cHi-C (**Fig. 1d**), and revealed a similar pattern of TADs, loops and stripes (**Fig. 1a** and **Extended Data Fig. 1a**). Comparing the contact frequency maps from hindlimb and forelimb, we see that the associations of *Pitx1* with *PelB, RA3, RA4*, and *Pen* are all more frequent in the hindlimb, where these enhancers contribute to *Pitx1* expression ^18–20^ (**Fig. 1c,f**), consistent with previous reports ^18^. These differences are also apparent when we examine the absolute distances instead of contact frequencies (**Extended Data Fig. 1d-f**). In contrast to these enhancer-promoter interactions, the interactions between *Pitx1* and the gene *Neurog1* are more frequent in forelimb, also consistent with previous cHi-C measurements ^18^. Although yet to be tested, this gene-gene interaction is likely mediated by Polycomb dependent long range looping ^29,30^, as both genes are enriched in Polycomb-associated H3K27me3 in forelimb, whereas in hindlimb, only *Neurog1* is H3K27me3 rich (**Fig. 1b**). The agreement between the two technically distinct assays testifies to the accuracy of the data. Additionally, the single-molecular nature of ORCA data allows us to robustly test the statistical significance of the difference across thousands of measurements (**Fig. 1f** and **Extended Data Fig. 1e**).

Surprisingly, while several cross-TAD interactions are more frequent in hindlimb, the TAD borders are also stronger in hindlimb compared to forelimb. For example, while *RA4* shows a higher frequency of proximity to *Pitx1* in hindlimb cells compared to forelimb cells, the TAD boundary B1 (next to RA3) is stronger, and the rest of the TAD containing RA4 (though not RA4 itself) is on average farther away (**Extended Data Fig. 1d**) and experiences fewer contacts with the TAD containing *Pitx1* (**Fig. 1c**, upper dashed box and middle circle). Similarly, the TAD containing the enhancer, *Pen*, is also on average farther away from and better insulated from the TAD containing *Pitx1*, even though *Pen* and *Pitx1* are on average closer (**Extended Data Fig. 1d**) and in more frequent contact (**Fig. 1c**, lower dashed box and circle). Since the measurements are made without the need for matrix balancing to address amplification biases, are imaged on the same slide, and replicated across independent experiments (**Extended Data Fig. 2**), we interpret these data to reflect true changes in the absolute interaction and not artifacts of normalization. We thus have a paradox: stronger TAD borders, yet more frequent enhancer-promoter contacts across borders.

We hypothesized this could occur through three plausible models (**Fig. 2a**). In the first model, “Merge”, the TADs merge in a subset of chromatins, for example due to the stochastic release of boundary factors. This would allow contact between enhancer and promoter in this subset, even though at the population level the boundaries would still be detected due to the contribution of other chromatins. In the second model, “Stack”, the TADs generally remain intact, but their boundaries stack around a central hub at which the boundary-proximal enhancers and promoters meet. Finally, it is possible that enhancer and promoter interact without the TADs merging or stacking, which we call “Loop out”. For example, the promoter and enhancer could be decompact enough to leave their respective TADs and meet outside the domain. These three models, as they stipulate hypotheses about how one structural aspect influences another in the very same molecules, cannot be resolved by Hi-C or other bulk methods that are restricted to measuring average pairwise interactions. ORCA, however, with its single-chromatin resolution, is well-suited for analyzing these models.

**Fig. 2.**
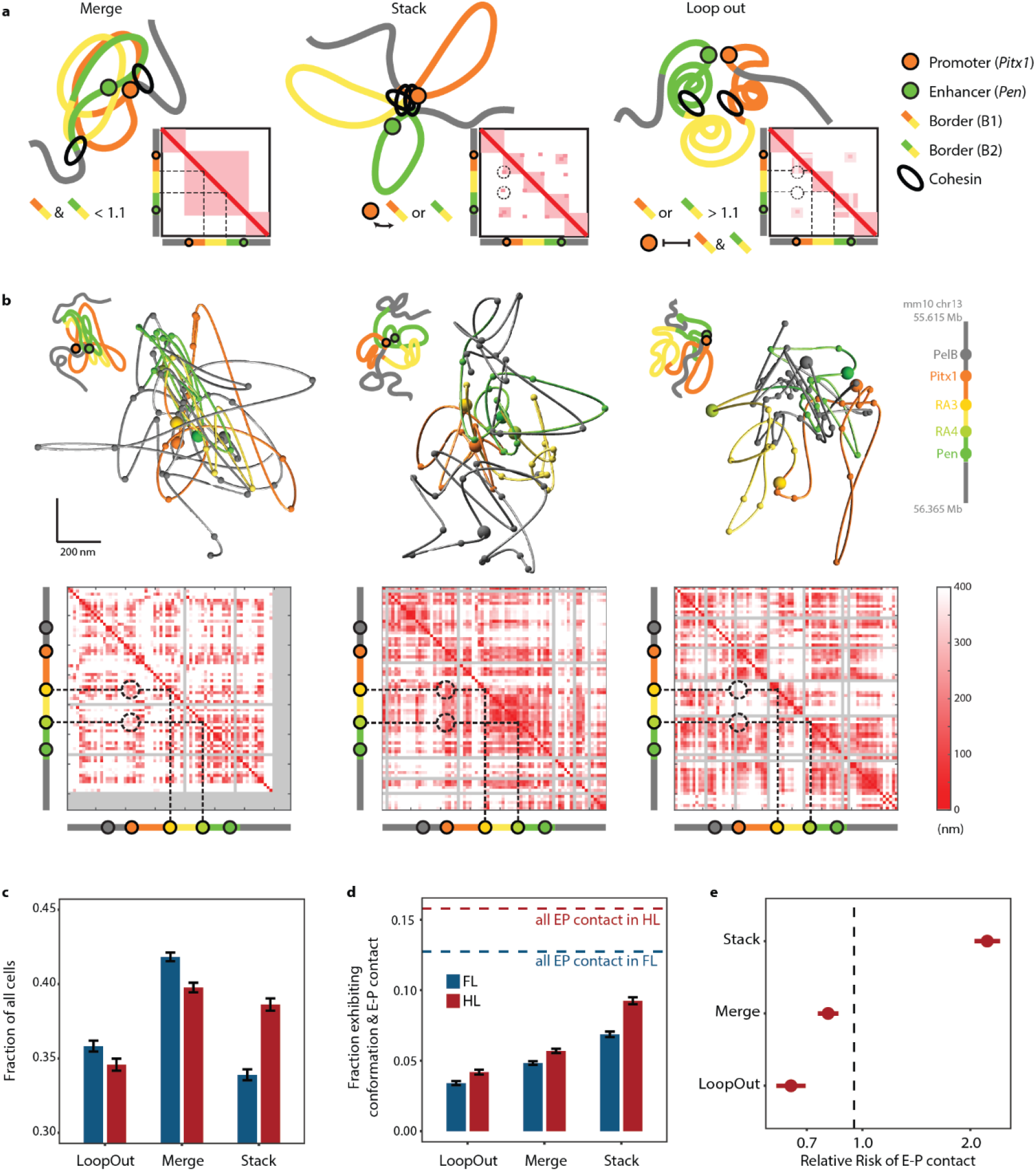
Testing three hypotheses regarding cross boundary enhancer promoter interaction. **a,** Schematic representation of the three models, each showing a cartoon chromatin trace, an idealized distance matrix at lower right, and the defining criteria at lower left. “Merge” is characterized by the weakening of boundaries B1 (orange-yellow) and B2 (yellow-green) between the enhancer and the promoter, emphasized by the dotted lines. “Stack” is characterized by promoter contacting intermediate boundaries B1 or B2, emphasized by the dotted circles. “Loop out” chromatins are those that neither merge nor stack (clear boundaries along the dotted lines and no contact in the dotted circles). **b,** Representative chromatin traces for each model. The top row shows reconstructed chromatin traces along with simplified cartoons that highlights the positions of enhancer, promoter and boundaries. The bottom row shows single-trace distance matrices. Gray rows and columns indicate missing data. The dotted lines mark the boundaries B1 and B2. The dotted circles mark the contacts between *Pitx1* and B1 and B2. Note how the contacts are present in “Stack” but not “LoopOut”. **c,** Fraction of all molecules exhibiting configurations matching each model in forelimb (blue) and hindlimb (red). Error bars represent standard deviations. **d,** Fraction of molecules where *Pitx1-Pen* interact exhibiting configurations matching each hypothesis in forelimb (blue) and hindlimb (red). Error bars represent standard deviations. Dashed lines indicate the fraction of all molecules exhibiting *Pitx1-Pen* interaction. **e,** Relative risk of having *Pitx1-Pen* contact for hindlimb molecule configurations matching each model. Bar indicates 95% confidence interval. X-axis is in log scale.

To compare the three models against the many individual molecules visualized with ORCA, we define criteria to determine if each chromatin trace fits Merge, Stack, or Loop Out configuration (**Fig. 2b** and **Extended Data Fig. 3**). We then analyze whether the chromatin traces fitting a particular model are more likely to have *Pitx1-Pen* contact, and whether such traces are more abundant in hindlimb versus forelimb cells. We define “Merge” chromatin traces as those having an insulation score of less than 1.1 (<10% difference in intra- vs. inter-domain median pairwise distance, after correction for linear distance) for both TAD boundaries between *Pitx1* and the *Pen* enhancer (B1 and B2 (**Fig. 1a**)), “Stack” traces as those with *Pitx1* contacting either B1 or B2, and “Loop out” traces as those not belonging to either “Merge” or “Stack”. For each model, there are numerous chromatin molecules consistent with its configuration in both forelimb and hindlimb (**Fig. 2c**). Notably, “Stack” is the only configuration that is more abundant in hindlimb than in forelimb. Moreover, among molecules with *Pitx1* contacting *Pen*, about 55% can be accounted for by “Stack”, highest among the three models (**Fig. 2d**). In addition, the difference in fraction of *Pitx1-Pen* contacting molecules between forelimb and hindlimb is most pronounced among “Stack” molecules. When assessing the relative risk, an indicator of the probability of finding *Pitx1-Pen* contact given one of the three configurations, “Stack” again is the only one with a value > 1, indicating that chromatin traces in the “Stack” configuration are (in this case, > 2x) more likely to have *Pitx1* contacting *Pen* compared to molecules not in “Stack” configuration (**Fig. 2e**). These analyses show that “Stack” is the most plausible mechanism for hindlimb-enriched interaction between *Pitx1* and *Pen*. Noticeably, despite nearly all hindlimb cells expressing *Pitx1*, only ~10% of them exhibit a “Stack” configuration where *Pen* and *Pitx1* are in contact (**Fig. 2d**), suggesting that the hub organization in “Stack”, similar to the formation of TADs, is a dynamic conformation instead of a static structure.

Thus, the data indicate that while all three mechanisms, “Loop out”, “Merge”, and “Stack”, contribute to the overall contact frequency observed in hindlimb, “Stack” is the primary conformation the chromosomal domain takes for *Pitx1* to bypass multiple TAD borders and make preferential contact with its distal enhancer, *Pen*. Such preferential contact was shown to be required for hindlimb specific up-regulation of *Pitx1* and avoidance of ectopic forelimb expression ^18^. Transient boundary loss or looping out across the domain occurs, but they do not enhance the probability of E-P contact, as their relative risks < 1, nor show much difference in frequency between hindlimb and forelimb.

The “Stack” model also helps explain another feature commonly observed in both Hi-C and ORCA data: long stripes in contact frequency maps. A stripe indicates that a genomic position has higher contact frequencies with almost all positions in the neighboring genomic region than other positions in that region have with each other. In **Fig. 1a**, for example, one of the stripes emanates from *Pitx1* and extends through almost the entire probed chromosomal domain. Hi-C analyses have found similar stripe patterns throughout the genome, and in many cases the stripes also connect enhancers and promoters, prompting new efforts to understand their molecular origins and new models of enhancer - promoter communication ^31–34^. Based on Hi-C analyses and polymer modeling, it has been proposed that stripes arise from the dynamic activity of cohesin molecules that load at an enhancer or promoter which is adjacent to a CTCF-site and reel-in the cognate enhancer/promoter through loop extrusion in the direction not blocked by CTCF ^31^. By sliding the loading site across the intervening DNA in this fashion, a stripe arises in the population map (**Fig 3a**). The “Stack” model provides an alternative explanation for the stripes. With elements next to CTCF sites stacked together at a central hub, these elements are simultaneously closer to all other elements in the domain (out in the intervening loops) than those other elements are to one another (**Extended Data Fig. 4**). The centrally located boundary elements are thus more likely to contact any element out in the loops through stochastic fluctuations of the polymer (**Extended Data Fig. 4c**).

**Fig. 3.**
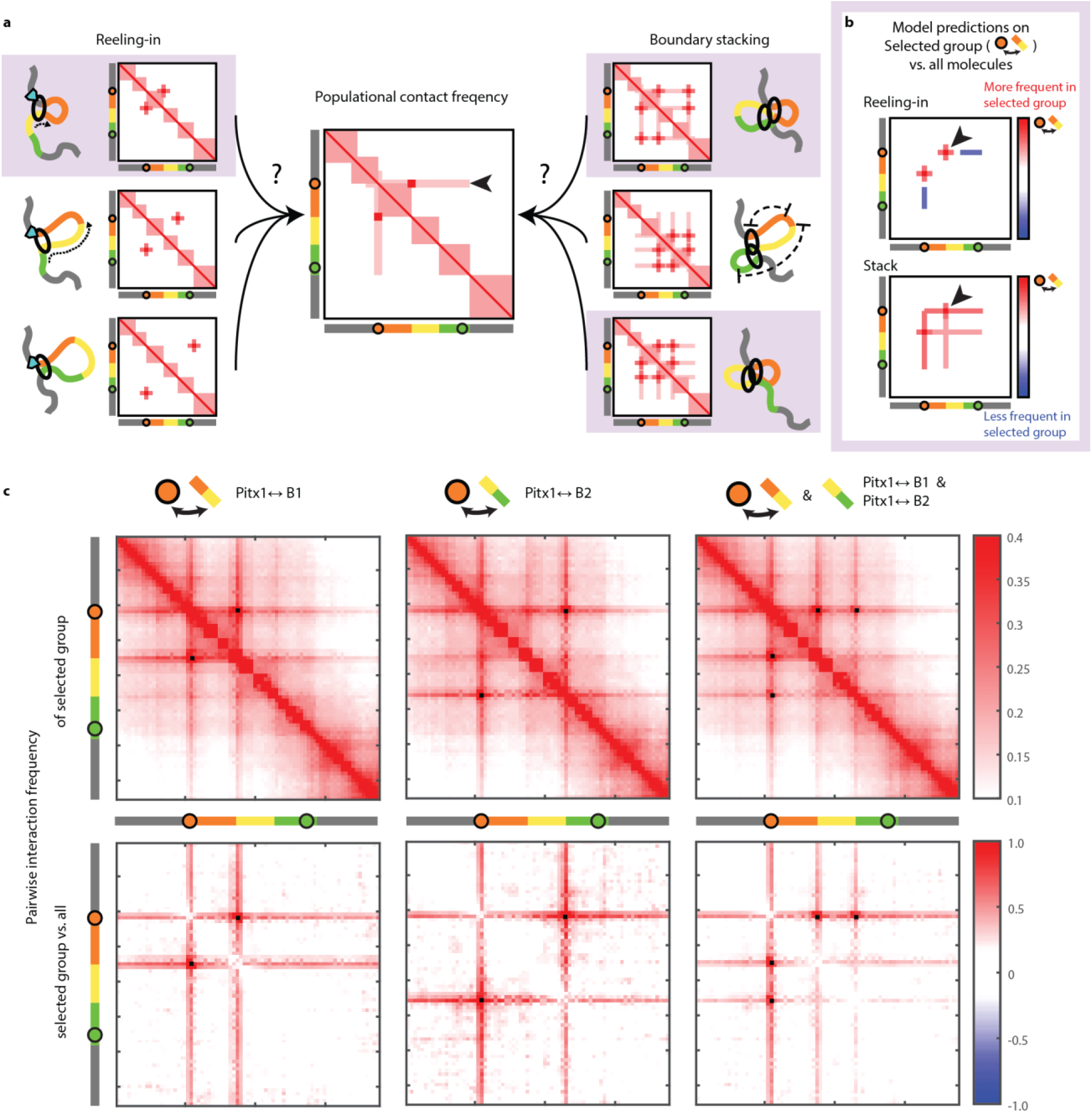
“Stripes” in single molecule data. **a,** Two models for the origin of “stripes”. (Center) Population-average pairwise contact frequency of the *Pitx1* locus in hindlimbs shows stripe patterns (arrowhead, refer to **Fig. 1a** for actual data.) (Left) Schematic representation of how stripes result through a “reeling-in” mechanism. The anchor at the CTCF site (cyan wedge) on the top, due to directional loop extrusion, interacts more frequently with the colored region compared to any other positions within the same region. Individual chromatin traces are each responsible for a dot along the stripe. (Right) Schematic representation of how stripes result from a “Stack” hub organization. Strong CTCF sites are collected in the central hub in 3D conformation. Due to its central location, the average distance from the hub to any position is shorter compared to pairwise distances between non-hub positions, as illustrated in the center-right panel. **b,** Testing the two models with single-molecule data. When comparing with all molecules, a selected group whose *Pitx1* and B1 are interacting (eg. shaded molecules in **a**) will by definition have a stronger “loop” between the said two positions, regardless of the model (arrowheads). However, “reeling-in” model predicts that other positions along the stripe would have less interaction compared to the population (blue lines), whereas “Stack” model predicts the opposite. **c,** (Top) Pairwise interaction of selected molecules on which *Pitx1* is contacting B1 (left), B2 (center) or both (right). (Bottom) Comparison of selected groups with the whole population, shown as their log2 ratio. The black pixels indicate the interactions used to select the group.

While both scenarios result in stripes at the populational level, they predict different behaviors at the single-molecular level. The “reeling-in” model predicts that for most molecules, *Pitx1* contacts no more than 1 position along the stripe: the stripe arises from capturing different chromatins in each step along the scan. Therefore, when examining chromatin traces where *Pitx1* contacts, for example, B1, *Pitx1* should not also be contacting the region between B2 and *Pen*, and the stripe should not be observed in each individual molecule. However, as seen in **Fig. 3c**, our single-molecule data shows that stripe is not only present, but stronger for molecules where *Pitx1* is interacting with either B1 or B2. The “Stack” model explains this phenomenon well: By conditioning on *Pitx1*-intermediate boundary interaction, we enrich for molecules with hub formation, a configuration where the boundaries are closer to all intervening regions. It is worth clarifying that we are not claiming “reeling-in” does not occur. To the contrary, it is likely through “reeling-in” that multiple domain boundaries stack together to form the “Stack” organization.

However, examining individual molecules suggests that stripes at the populational level are not merely an aggregate of single loops. Instead, “Stack” hub organization is seen in many contributing molecules, suggesting that it is indispensable for interpreting the underlying dynamics of these stripes. We observed similar aggregates of loop hubs in individual traces producing stripes in the population maps for the *SOX9* locus in human cranial neural crest cells, though in that case the stripes did not extend beyond TAD boundaries ^35^.

We next asked, what molecular mechanisms facilitate the formation of the “Stack” conformation, and what molecular conditions differ between hindlimb and forelimb to bring this about in a tissue-specific manner? To answer these mechanistic questions, we turned to physical polymer modeling. Using the polymer modeling framework from open2c ^36^, derived from prior work modeling loop-extrusion dynamics in chromatin ^37,38^, we simulate the dynamics of a chromosomal domain with 4 extrusion blockers, representing *Pitx1* (promoter), B1, B2 and *Pen* (enhancer), respectively, each of which correspond to TAD-borders marked by CTCF (**Fig. 1a**). We use these simulations to explore interactions among sequentially arrayed border elements under distinct regimes of loop extrusion.

In our simulation, increasing CTCF occupancy by modeling not only moderately strengthened the TAD boundaries as expected, it also strengthened the cross-TAD contact frequency between the *Pen* enhancer and the *Pitx1* promoter (**Fig. 4e**). Examination of polymer structures with E-P contact showed frequent involvement of the intervening boundaries B1 and B2, consistent with the “Stack” organization seen for the *Pitx1* domain. Plotting the positions of the cohesin molecules reveals the presence of multi-cohesin bridges connecting the promoter to B1 to B2 to the enhancer (**Fig. 4a**). Although not all E-P contacts involved the intermediate boundary interactions, indicating that sometimes “Merge” or “Loop out” rather than “Stack” allows contact, “Stack” configuration accounts for most of the increase in contact upon increased CTCF occupancy (**Fig. 4f,g**). Similarly, the relative risk for observing E-P contact given interaction with intermediate boundaries is significantly greater than 1, whereas the relative risk given either “Merge” or “Loop out” configurations is not (**Fig. 4h**).

**Fig. 4.**
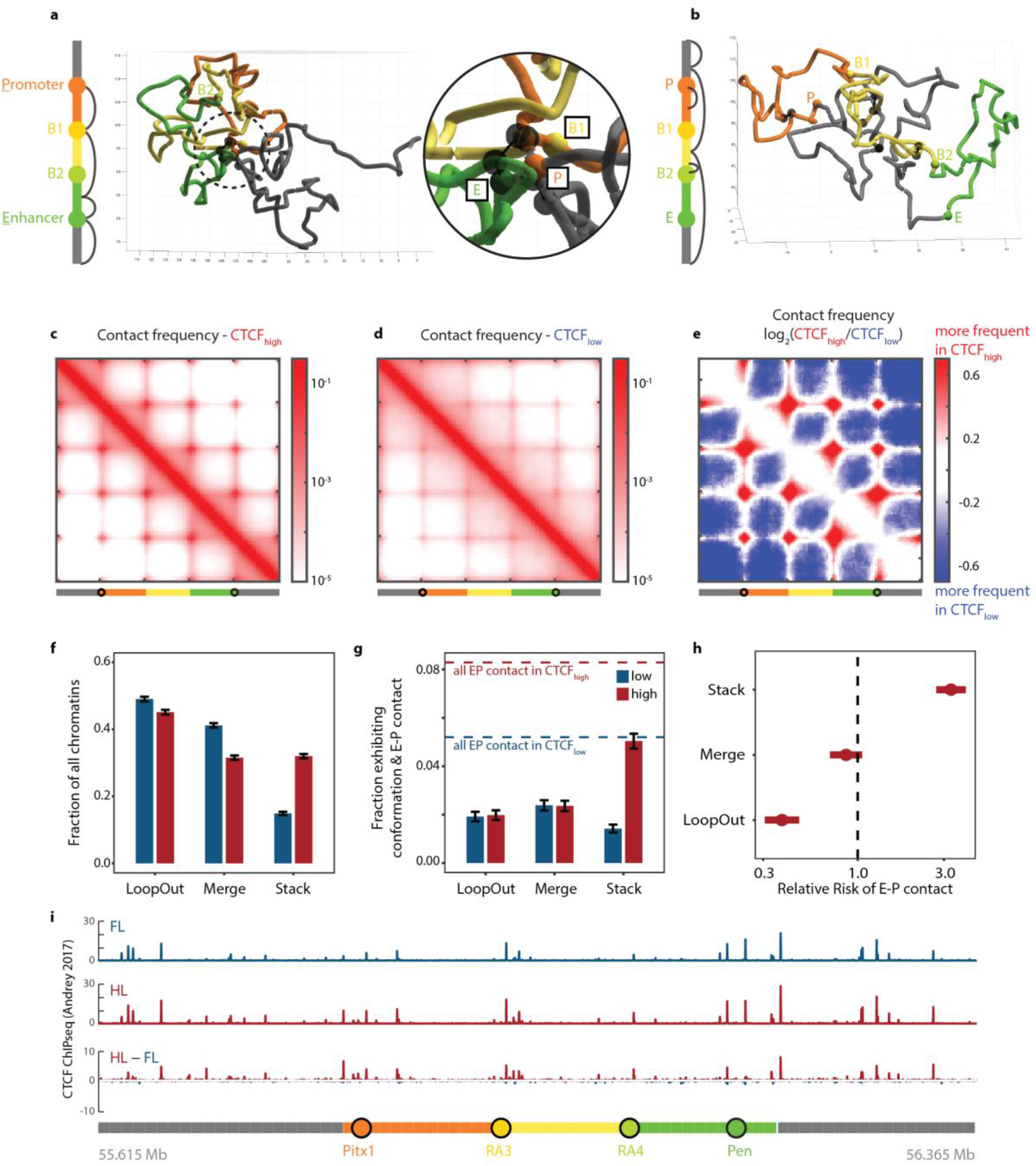
Modeling effects of increasing CTCF activity. **a, b,** Snapshots of individual simulations. The bars on the left depict the polymer used in simulation straightened, with a color scheme parallel to that used for the actual *Pitx1* chromosomal domain. The colored circles are CTCF sites. Black arcs represent positions brought together by cohesin molecules in that snapshot. **a,** A polymer with high CTCF binding rate and thus more cohesin (transparent black dumbbells) arrested at each CTCF site. The inset shows a zoomed-in view of the circled area, where 3 CTCF sites, the promoter (P), enhancer (E) and intermediate boundary B1, are brought together by arrested cohesins stalled between them in a bridge. **b,** Simulation with low CTCF concentration. **c,** Contact frequency at high CTCF. **d,** Contact frequency at low CTCF. **e,** log2 ratio of data in **c** and **d**. **f,** Fraction of all simulated polymers exhibiting configurations matching each model in low CTCF (blue) and high CTCF (red) conditions. Error bars represent standard deviations. **g,** Fraction of all simulated polymers that have E-P interaction exhibiting configurations matching each model in low CTCF (blue) and high CTCF (red) conditions. Error bars represent standard deviations. Dotted lines represent the fraction of all polymers exhibiting E-P interaction. **h,** Relative risk of having E-P contact for polymers under high CTCF condition while exhibiting configurations matching a model. Bar indicates 95% confidence interval. **i,** Comparison of CTCF ChIP-seq peak heights ^21^ in hindlimb (red) vs. forelimb (blue) at E10.5.

To explore this effect more generally, we do not insist on matching the specific organization of the *Pitx1* region with respect to the distances between border elements, their relative orientation or strengths. However, additional simulations, in which we manually tuned these parameters to give a more *Pitx1-like* structure to that observed experimentally (Pearson’s R=0.93), recapitulated the effect of boundary strengthening in increasing boundary bypass (**Extended Data Fig. 5**).

As a test of this hypothesis, we examined the binding of CTCF across the *Pitx1* domain in hindlimb vs. forelimb in published ChIP-seq data ^21^. Interestingly, CTCF peak heights were higher at *Pitx1*, B1, B2 and B3 in hindlimb than forelimb, even though background binding levels between the peaks were similar (**Fig. 4i**). This suggests that differences in CTCF binding to peak regions could explain the distinct structural organization of the *Pitx1* domain in hindlimbs, which allows sufficient interaction for activation by *Pen* specifically in hindlimb.

To see if the Stack model applies more broadly outside of the *Pitx1* domain, we explore published datasets to examine the relationship between enhancer-promoter (E-P) pairs and TAD boundaries. If the Stack organization is a major mechanism for border-crossing interactions, we would expect that cross-TAD interacting *cis*-elements are more likely to be closer to boundaries than intra-TAD interacting *cis*-elements, whose adjacency to boundary matter less since there are no borders to cross. For E-P pairs, we take advantage of the Transcribed Enhancer Atlas from the FANTOM 5 project, where enhancer-promoter pairs were mapped using the correlation of mRNA and eRNA transcription in capped analysis of gene expression (CAGE) results across diverse human cell types ^39^. For boundary calling, we use the Hi-C data obtained from GM12878 cell line ^40^ (**Fig. 5a**). We deliberately choose these datasets despite the lack of recency. Rao *et al*. remain one of the highest-resolved human Hi-C maps and demonstrated a high degree of conservation of boundaries across cell types. The FANTOM 5 E-P pairs were determined in a TAD-agnostic way and a substantial number of them were > 100 kb apart. Most more recent E-P prediction datasets took TAD boundaries into consideration and/or uncovered relatively few E-P pairs >100 kb apart, possibly due to limitations of cell-type choice and assay sensitivity ^41,42^, rendering them inappropriate for our purpose.

**Fig. 5.**
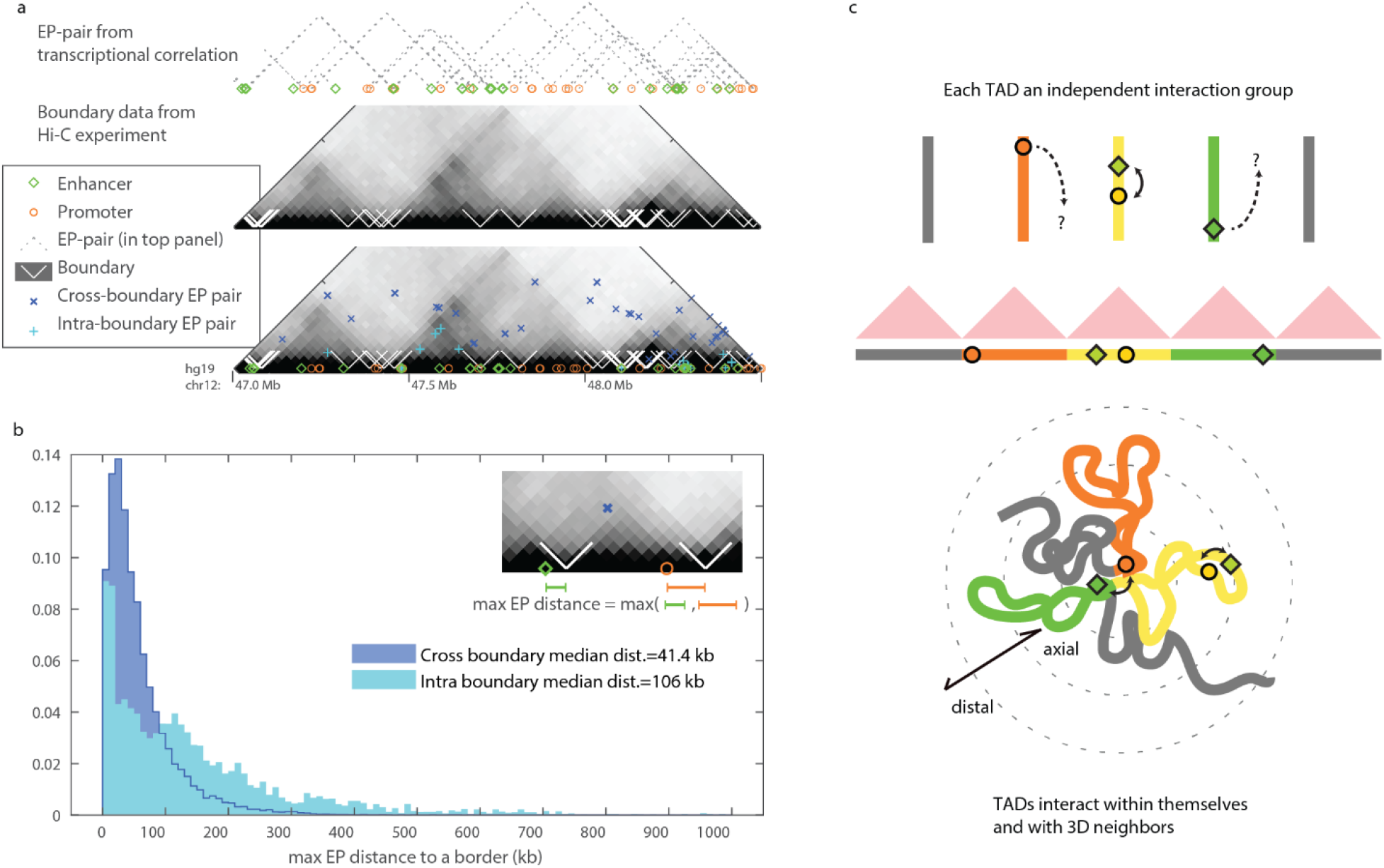
Genome-wide analysis of cross-TAD enhancer-promoter interactions. **a, b, Boundary-proximity is enriched for E-P pairs that interact across boundaries. a,** Combining information of E-P pairs across the genome ^39^ and boundary information from Hi-C data ^40^, we categorized each E-P pair as cross-boundary or intra-boundary. **b,** Histogram of maximum distance to a border for E-P pairs that are >100 kb apart. **c, New perspective on the role of chromatin boundaries.** (Top) Conventional models consider each TAD as largely independent and do not account for cross-TAD interactions. (Bottom) Stack model adds in the 3D relationship between TADs and introduces the dimension of axial - distal position for each element relative to the whole molecule.

We focus on long-range E-P interactions (>100 kb, ~60% of the 1e5 E-P pairs in the dataset), and intersect these with TAD border calls from Hi-C. We find many of these long-range interactions cross a TAD border. To see if border-stacking could play a role in a substantial fraction of these cross-border interactions, we compute the largest distance to the nearest border for both E and P in boundary-crossing pairs and for non-boundary-crossing pairs. We find crossing pairs are significantly more likely to be near a border (*p*=2e-308, Mann-Whitney U and KS-test) than non-crossing pairs, with a median distance of 40 kb from the nearest border for both E and P, rather than over 100 kb (**Fig. 5b**). This statistical trend is consistent with border stacking facilitating border bypass for a substantial number of E-P interactions across the human genome.

## Discussion

Here we set out to understand the physical and molecular mechanisms which allow some enhancers to regulate genes across TAD boundaries. Combining imaging and molecular dynamic modeling, we arrived at a model in which strengthening the loop-extrusion-blocking behavior of border elements increases the probability that they will be collected together at a central hub by arresting consecutive cohesin-mediated loops. Enhancers and promoters proximal to these borders thus also gather at the hub, where they interact with elevated frequency, whereas enhancers and promoters more distal from the border are neither intermeshed by extrusion nor collected at hubs. This model provides a mechanistic explanation of why some enhancers can act across TAD borders more easily than others, and a predictive basis on which to identify border-crossing enhancer candidates.

The picture of the Stack organization (e.g. **Fig. 5c**) has a superficial resemblance to earlier models of multi-enhancer-gene hubs, sometimes described as rosettes ^43^. Notable examples include interactions between the protocadherin alpha genes ^44^, the *Tcr* locus in thymocytes ^45^ and enhancer “archipelagos” near the *Hoxd* genes ^46^. However, they differ from the organization proposed here both in the molecular evidence supporting the model and in the way the structure was interpreted for gene expression. Rosette structures were proposed on the basis of 3C (and in some cases later 4C experiments), which showed preferential ligations among the genes and enhancers. As these methods measure only pairwise interaction, the proposed multi-way hub organization is by nature speculative. Entirely distinct conclusions were drawn from the inferred structure as well. The cell-type specific nature of enhancer-promoter interaction was emphasized, but no question of border and border bypass existed for these loci, nor was any speculation of boundary effects a component of the models. As Hi-C increasingly displaced 3C approaches and provided more ways to correct for the accessibility bias at enhancers and promoters ^47^, the enhancer rosette model at these loci has given way to a TAD interpretation ^48–50^. In investigating how a key developmental enhancer might bypass these TAD borders, we have presented direct data for specific multi-way interactions. Rather than functioning as enhancer hubs, these multiway interactions allow a 3D geometry that preserves boundary function for border-distal elements while enabling boundary-bypass for border-proximal elements.

In our model, the cell-type specific activation of *Pitx1* gene by the distant *Pen*-enhancer can be explained by the strengthening of extrusion blocking borders in hindlimb relative to forelimb. Experimental data indicates that one of the most notable increases in CTCF border binding occurs at the *Pitx1* promoter itself, with a corresponding increase in Rad21 detection (**Fig. 1b & Fig. 4i**). The strengthening of CTCF binding, especially near the *Pitx1* promoter, increases the frequency with which *Pitx1* joins enhancers at the hub. Such differential CTCF binding could be achieved through several different mechanisms. A simple hypothesis that developing hindlimb cells express more CTCF finds no support in published data surveyed so far, including results from mice ^21^, bat ^51^ and chicken ^52^. Other more probable hypotheses include tissue-specific expression of cofactors that either recruit CTCF to the *Pitx1* locus or open up binding sites for CTCF. Interference of CTCF binding through DNA methylation of CTCF binding sites could also play an important regulatory role ^53,54^. Although we are not aware of any methylomic dataset regarding forelimbs and hindlimbs of tetrapods, a previous study showed differential methylation of a transgene along the anterior-posterior axis in developing mouse embryos ^55^. Specifically, methylation level is higher in the forelimbs compared to hindlimbs, resulting in lower expression of the transgene in the forelimbs. If the methylation difference is in the same direction at the *Pitx1* locus, it would help explain the forelimb-hindlimb differences in CTCF ChIP-seq binding, chromatin conformation, and *Pen-Pitx1* contact frequency.

Our model provides an explanation for some other long-standing puzzles in developmental *cis*-regulation. From flies to mammals, genes in the Hox family are controlled by distal enhancers which act across CTCF-marked TAD boundaries ^10,22,56–62^. The *Hoxd genes* are regulated in the vertebrate limb by a series of enhancer “islands” scattered throughout the 1 Mb upstream region ^46,50^. Many of these islands, as well as the genes they regulate, are closely linked to CTCF borders ^13^. Similarly in human hematopoietic stem cells, enhancers of anterior *Hoxa* genes, located 1.3 Mb apart from their promoters and separated by TAD borders, are themselves juxtaposed to a strong TAD border ^11,63^. In Drosophila, the promoter of the Hox gene *Abd-B* sits near a boundary and communicates across intervening borders to its distal enhancers *iab-7* and *iab-5* ^10^, as does the Hox gene *Scr*, whose enhancers also bypass the intervening gene *ftz* ^62^. Several of these boundaries (a.k.a. insulators) not only coincide with TAD borders but have been shown to functionally block enhancer-promoter communication in a transgene context and prevent ectopic activation at native loci. Leading geneticists termed this cross-border regulation puzzle “the boundary paradox” ^10^. Our model predicts that these borders, rather than inhibit or attenuate, actually facilitate action of the distal enhancers through the preferential stacking of the borders themselves. These examples of border-crossing, boundary-associated enhancers of a few major developmental control genes, along with our genome-wide analysis on the relationship between tens of thousands of inferred E-P pairs and their distances to TAD boundaries, suggest that the border stacking model proposed from *Pitx1* could be informative more generally. Such generality provides a new perspective on the contexts in which insulators will be most and least effective. Adding an insulator between an enhancer and a promoter already anchored by CTCF boundaries may have more subtle effects on contact interactions than when either the enhancer, promoter or both are not closely associated with TAD borders. This may contribute to the surprising recent findings that a variety of previously characterized insulators provide little attenuation between *Sox2* and its distal *SCR* enhancer ^64,65^.

Domain boundaries and the process of loop extrusion have long been two of the fundamental concepts by which we contemplate the dynamics of chromatin organization. Despite their billing as features of “3D genome organization”, TADs and their boundaries are often thought-of primarily in a quasi-1D manner. In the past, boundaries were regarded as barriers to genomic cross talk: elements only interact with other elements in the same TAD, and each TAD functioned independently from each other (**Fig. 5c**, top panel). In this view the case of *Pen* and other border crossing elements appear to be unexplained exceptions. Our 3D image data adds an additional dimension to this quasi-1D model. While multiple borders cluster together in the Stack organization, border-proximal enhancers and promoters are brought into proximity, even if separated by multiple TADs. Meanwhile, border distal regions of the TAD extrude radially from this hub and are not intermixed, allowing the border function to persist (**Fig. 5c**). Moreover, this border stacking organization is consistent with the loop-extrusion model, as long as cohesin is sufficiently dense on chromatin for multiple cohesins to arrest at opposite sides of an intermediate border, facilitating the formation of a stack. Our model adds a new framework to help us reconcile loop extrusion and TAD borders with previously paradoxical examples of long-range *cis*-regulation, including border bypass and gene skipping behavior, bringing us a step closer to a predictive understanding of genome function.

## Acknowledgements

This work was supported in part by a New Innovator’s Award from the NIH, a Beckman Young Investigator Award, and a Packard Foundation Award (ANB); a predoctoral fellowship from the Ministry of Education of Taiwan (TH); and an HHMI Investigator position (DMK).

## Material and Methods

### Animal procedures

All animal procedures followed guidelines approved by Stanford University’s Administrative Panel on Laboratory Animal Care (APLAC).

### Mouse limb bud section preparation

C57Bl/6 embryos were harvested at E11.5 or E12.5. Pregnancy was timed by checking vaginal plugs and the developmental stage is confirmed by embryo morphology. The pregnant mouse was euthanized with CO_2_ inhalation for 8 minutes followed by cervical dislocation. Limb buds were dissected in PBS and fixed in 4% PFA in PBS at 4°C for 2 hours. (Fixation reduces background signals.) Afterwards limb buds were washed in PBS at 4°C for 5 minutes 3 times and then individually oriented in OCT blocks, frozen in liquid nitrogen and preserved in −80°C. For sectioning, forelimb-bud and hindlimb-bud of the same embryo from the same side (right or left) were sectioned together. The OCT blocks were placed at −20°C for 30 minutes to equilibrate. Limb buds were sectioned from the proximal end along the proximal - distal axis and parallel to the anterior - posterior axis with 8 μm thickness. Sections of forelimb-bud and hindlimb-bud were placed on the same slide, which was coated with chromium–gelatin.

For additional details, please refer to “Embryo cryosectioning” (steps 69 - 75) of ^23^. Note that for all our samples, we proceeded to probe hybridizing immediately without storing the slides in the fridge or freezer.

### ORCA experiment

We closely followed the method comprehensively described in ^23^.

### Probe Design

We tiled the mouse *Pitx1* chromosomal domain, mm10 chr13:55,615,000-56,475,00, with 86 10-kb barcodes, each consisting of ~250 individual probes. Each probe is 80 bp long, with 20 bp complementary to fiducial probes, followed by 20 bp of barcode, and ended with 40 bp of unique target sequence. The probes were designed by the software described previously ^23^. Note that we only report results for the first 75 barcodes, as not all experiments used all 86 barcodes, and regions beyond barcode 75 were not crucial for the main conclusions in this work. The complete sequences of probes and their genomic targets can be found on the 4DN data portal. The probe design software can be found on https://github.com/BoettigerLab/ORCA-public.The probe library was ordered from CustomArray (now operated by Genscript) as an oligo pool.

### ORCA hybridization

We followed the procedure from ([Tracing DNA paths and RNA profiles in cultured cells and tissues with ORCA | Nature Protocols](https://www.nature.com/articles/s41596-020-00478-x)). See “Hybridization with DNA primary probes”, steps 147 - 166. Following are details the protocol did not specify or modifications for our experiment. For most steps, the slide was placed inside a plastic petri dish with a diameter of 6 cm, with buffers added by pouring from a 50 mL tube and removed by suction. During the 37°C incubation with RNaseA (step 152), the petri dish containing the slide was placed inside a humidifying chamber (a tip box with water inside). We used a 22 x 22 mm coverslip at steps 154 and 157. At step 156, our hybridization mixture was composed of 26 μL of hybridization no. 2 plus 3 uL of 500 ng/uL primary probe (29 uL in total).

### Imaging system setup

For the microscope setup, see ^22^. (Also the “Microscope setup” section in Introduction of ^23^.) All experiments were imaged on the system with the Ti2 body, which used a 1,536 x 1,536 (px) field of view. Briefly, it is composed of an IR-laser based autofocus system, a high performance 3D stage with 500 nm range piezo based z-stage (Ludl), and a custom fluidics system. Illumination is provided by a 561-nm solid-state laser (MPB 2RU-VFL-P-2000-560-B1R, for fiducial probes) and a 647-nm solid-state laser (MPB 2RU-VFL-P-2000-647-B1R, for readout probes). The microscope setup is available on the Micro-Meta: https://data.4dnucleome.org/microscope-configurations/33ea326f-a557-4457-becf-55bc860c4bdf/).

For the automated fluidics system setup, see the “Fluidics setup” section in Introduction of ^23^. Briefly, a custom built robotic system with a 3-axis CNC router engraver, buffer reservoirs and hybridization wells (96-well deep well plate) on the 3-axis stage, ETFE tubing, imaging chamber (FCS2, Bioptechs), a needle, and peristaltic pump (Gilson F155006). The needle moved between buffers of hybridization wells and flowed the liquid across the sample in the imaging chamber through the tubings with the pump.

### Imaging process

For the imaging process, see the “ORCA imaging” section in ^22^ and steps 167 - 197 of ^23^. Briefly, we used open-source software to control and coordinate the fluidic system and the microscope. First, 2x SSC buffer was flown into the sample chamber, and fields-of-view (FOV) were selected to cover either various parts or the whole limb section. Each FOV was then photo-bleached with both 561-nm and 647-nm at maximum intensity for 1 - 3 minutes.

For each readout barcode, we first flowed in 200–500 μL probe solution (25% ethylene carbonate and 2x SSC containing the readout probe and oligos complementary to the previous readout probe to peel off remaining probes) and incubated for 15 mins to hybridize. Then we flowed 1 mL of wash buffer (30% formamide in 2x SSC buffer) for 2 mins, followed by washing with 1 mL of 2x SSC buffer for 2 mins. At the imaging stage, we filled the chamber with imaging buffer (0.5 mg/ml glucose oxidase, 40 μg/ml catalase and 10% w/v glucose in 2x SSC). Each FOV was then sequentially imaged with 561-nm and 647-nm in z-stacks of 100-nm steps for a total of 100 steps. After all FOVs were imaged, 2x SSC was flown and each FOV was photobleached for 3 seconds by the 647-nm laser. This process was repeated until all barcodes was imaged. Depending on the number of FOVs, each round of imaging took ~1 - 2 hours.

### Analysis

Image processing was performed following steps 198 onward of ^23^. For each experiment, the resultant data are the 3D coordinates of each barcode of each detected chromatin. For each detected chromatin, its 3D coordinates can be converted into a pairwise distance matrix for ease of analysis.

### Masking of bad hybridizations

When presenting population-average level data, either contact frequencies or median pairwise distances, we mask the bad hybridizations and interpolate with adjacent values. Bad hybridizations, namely 16, 21, 26, 29, 30, 32, 33, 36, 39, 40, 44, 49, 53, 63, 65 (refer to **Fig. 1a** for their genomic positions), are determined by visual examination of the raw imaging data, where either weak signal or off-target bright dots are present. Many of these bad hybridizations are a result of some intrinsic properties of the readout probes. The same probes have yielded poor hybridization in ORCA experiments targeting completely different regions in different organisms.

### Normalization

The average 3D physical sizes of the probed region may vary among experiments due to variations in time a particular slide spent in different reagents, as well as how individual limbs were snipped and subjected to fixation. To address this batch effect when comparing between experiments, we normalized the median of all pairwise distances between probes 1 - 68 (common to all experiments) measured in a given limb in a given experiment.

### Insulation score (categorizing “Merge” molecules)

We defined the insulation score of a TAD boundary for a single molecule as the median cross-TAD pairwise distances divided by the median intra-TAD pairwise distances.

### Simulation

Polymer simulations were performed with the open source polychrom software ^36^ from open2c. Codes for performing and analyzing results, including the parameters used in each condition, are available on our Github repository. Briefly, the software constructs Langevin dynamic simulations of flexible polymers moving under thermal noise with user defined energy potentials to describe the dynamics of the polymer and molecular interactions among monomers, including the links produced by loop extrusion factors. The simulations use the freely distributed openMM framework ^66^ to provide GPU accelerated computation. For the generalized simulations in **Fig. 4**, the CTCF loading rates for each CTCF binding site are 0.18 and 0.7 for the low and high CTCF conditions, respectively. For the fine-tuned simulations in **Supp. Fig. 5**, the strength of each CTCF binding site is different, with the low CTCF condition having a 60% CTCF loading rate compared to that of the high CTCF condition.

### Data availability

The chromosome trace data has been converted to the NIH 4DN data standard, FOF-CT (FISH-omic Format, Chromosome Tracing) and can be accessed at 4DN data portal (https://data.4dnucleome.org/) with accession 4DNES4TC13IL. Simulation data are available on Zenodo with the following DOIs: 10.5281/zenodo.7566077, 10.5281/zenodo.7566087, 10.5281/zenodo.7566087, 10.5281/zenodo.7572059.

### Code availability

Scripts for analyses and figure generation are available in our github repository: https://github.com/BoettigerLab/ORCA-Pitx1-2022. Polymer simulations also require the simulation toolkit adapted from the open2c project, which is available here: https://github.com/BoettigerLab/polychrom. Probe design and image analysis software is available at https://github.com/BoettigerLab/ORCA-public.

## Extended Data Figures

**Extended Data Fig. 1.**
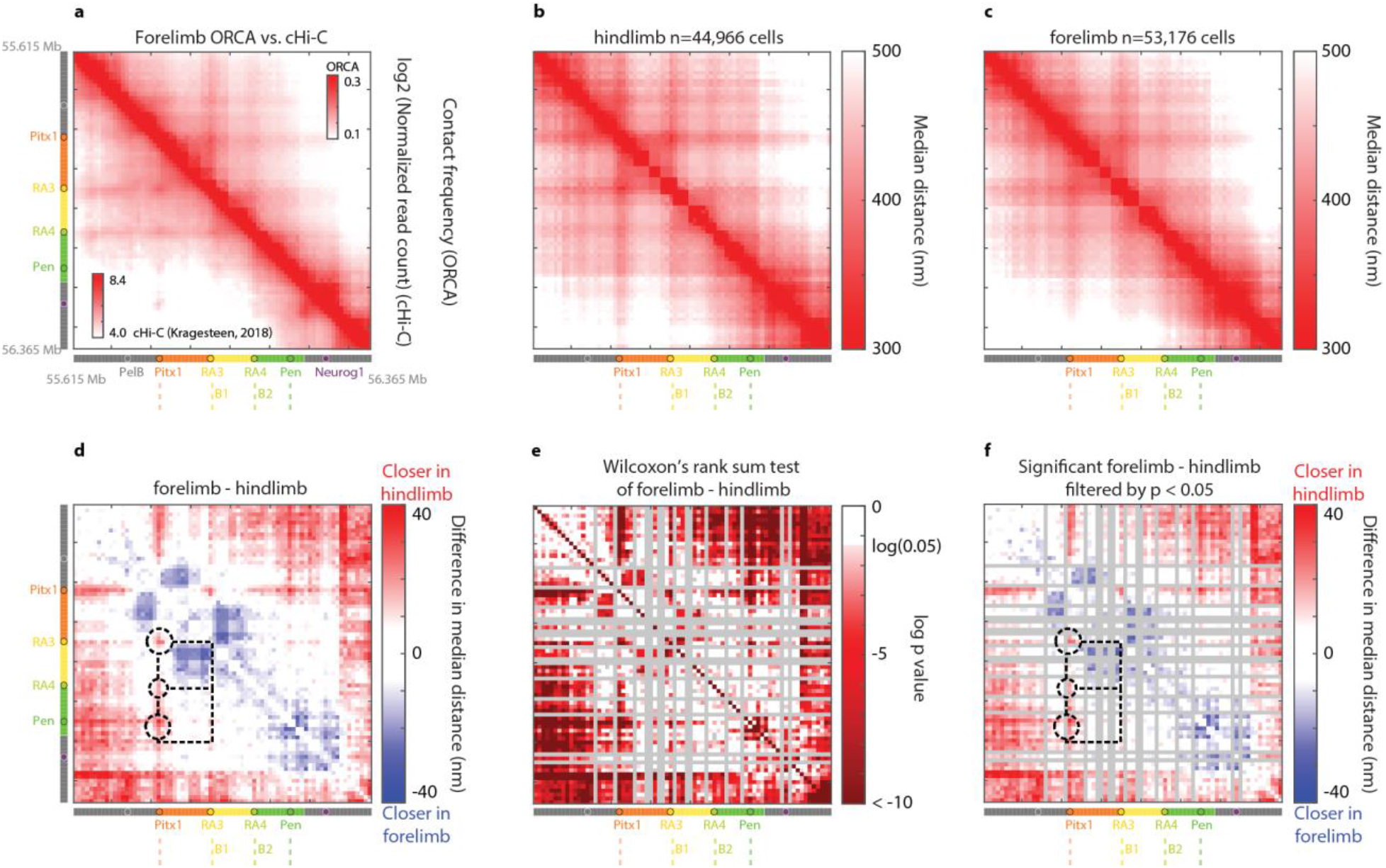
Forelimb contact frequency and median pairwise distance data. **a,** Chromatin conformation at *Pitx1* domain in forelimb cells as detected by ORCA (upper-right half, from 53,176 individual chromatin traces) and cHi-C ^18^ (lower-left half). **b, c,** ORCA median pairwise distance of the *Pitx1* chromosomal domain in hindlimb and forelimb cells, respectively. **d,** Difference in distance between forelimb and hindlimb cells. Red indicates that distance is smaller (closer) in hindlimb. **e,** Matrix of p-values for all distance differences shown in **d**, using Wilcoxon rank-sum test for each pairwise distance. The colorbar turns from white to pink at p = 0.05. Bad hybridizations (see Material and Methods) are masked gray. **f,** Significant (raw p-value < 0.05) distance differences. Non significant differences are masked white.

**Extended Data Fig. 2.**
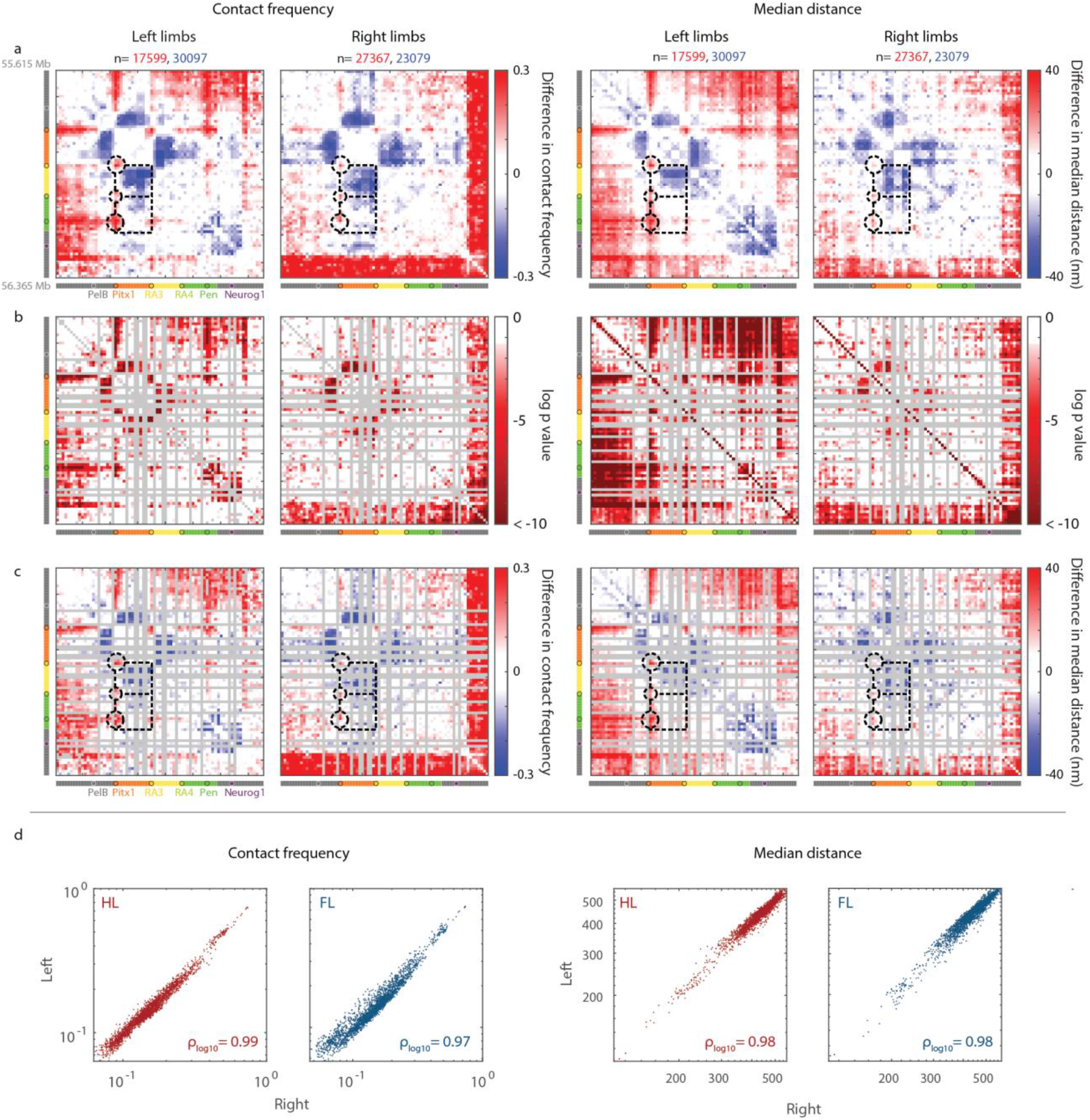
Concordance of results between independent ORCA experiments. The left two columns are the maps for forelimb-hindlimb differences in pairwise contact frequency from experiments using either left limbs or right limbs and independent hybridizations. The right two columns are the maps for forelimb-hindlimb differences in pairwise distances. The two numbers of n denote the number of chromatin traces observed for hindlimb and forelimb, respectively. **a,** The original maps. **b,** Matrix of p-values for all differences shown in **a**, using a chi-square test for each pairwise contact frequency and Wilcoxon rank-sum test for each pairwise distance. The colorbar turns from white to pink at p = 0.05. Bad hybridizations are masked gray. **c,** Significant (raw p-value < 0.05) differences. Non significant differences are masked white. For **a** and **c**, red indicates more contact or closer distance in hindlimb. Difference in contact frequency is shown as the log2 ratio of hindlimb to forelimb. Key features discussed in this paper, highlighted by the dashed boxes and circles, are consistent between experiments from left or right limbs, such as the hindlimb-enriched contacts between *Pitx1* promoter and its enhancers and forelimb-enriched contacts between orange TAD and yellow-green TAD. The “red borders” seen in the final hybridizations (68 - 75) of the right limb is likely an experimental artifact from the end of a run. It is only seen in experiment #1, and the other right limb experiment (#5, see **Supplementary Table 1**) does not have data for these hybridizations. **d,** Correlation between left and right limb experiments for pairwise contact frequencies (left panels) and median distances (right panels).

**Extended Data Fig. 3.**
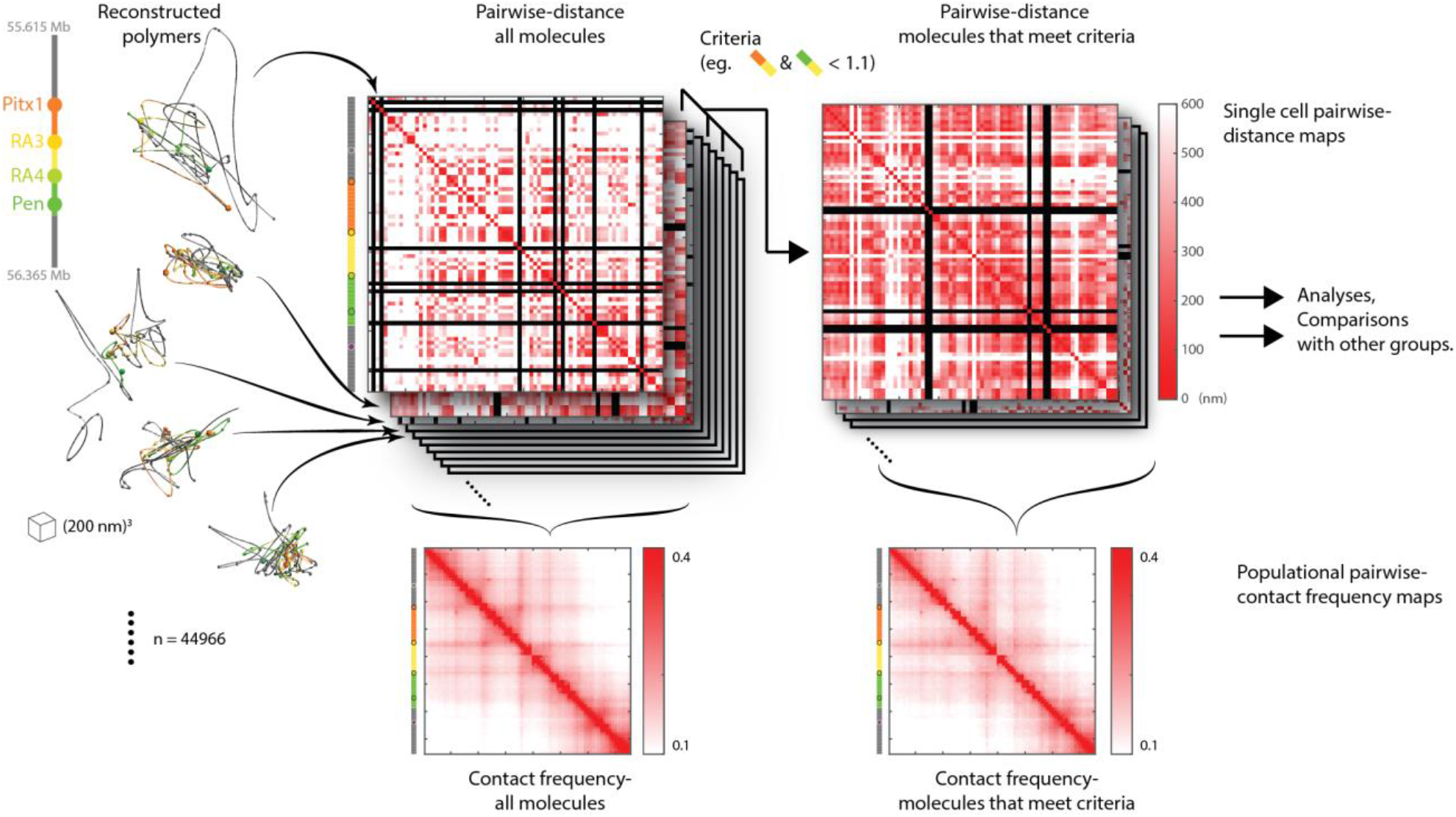
Selecting groups of molecules in ORCA data based on model criteria. This figure illustrates how, for example, “Merge” molecules or “Stack” molecules were selected from all molecules. (Left) First, the shape of individual *Pitx1* regulatory domains are reconstructed from raw ORCA data. (Center) Each molecule can be represented as a pairwise-distance map. (Right) Maps that match a certain criteria (in this example, weak boundary strengths on both B1 and B2) are selected into a group for further analysis. (See **Fig. 2 c-e**). (Bottom) For any groups of molecules, a populational representation can be obtained using a pairwise-contact frequency map, in which the value at (x, y) represents the percentage of molecules having the distance between positions x and y within 200 nm.

**Extended Data Fig. 4.**
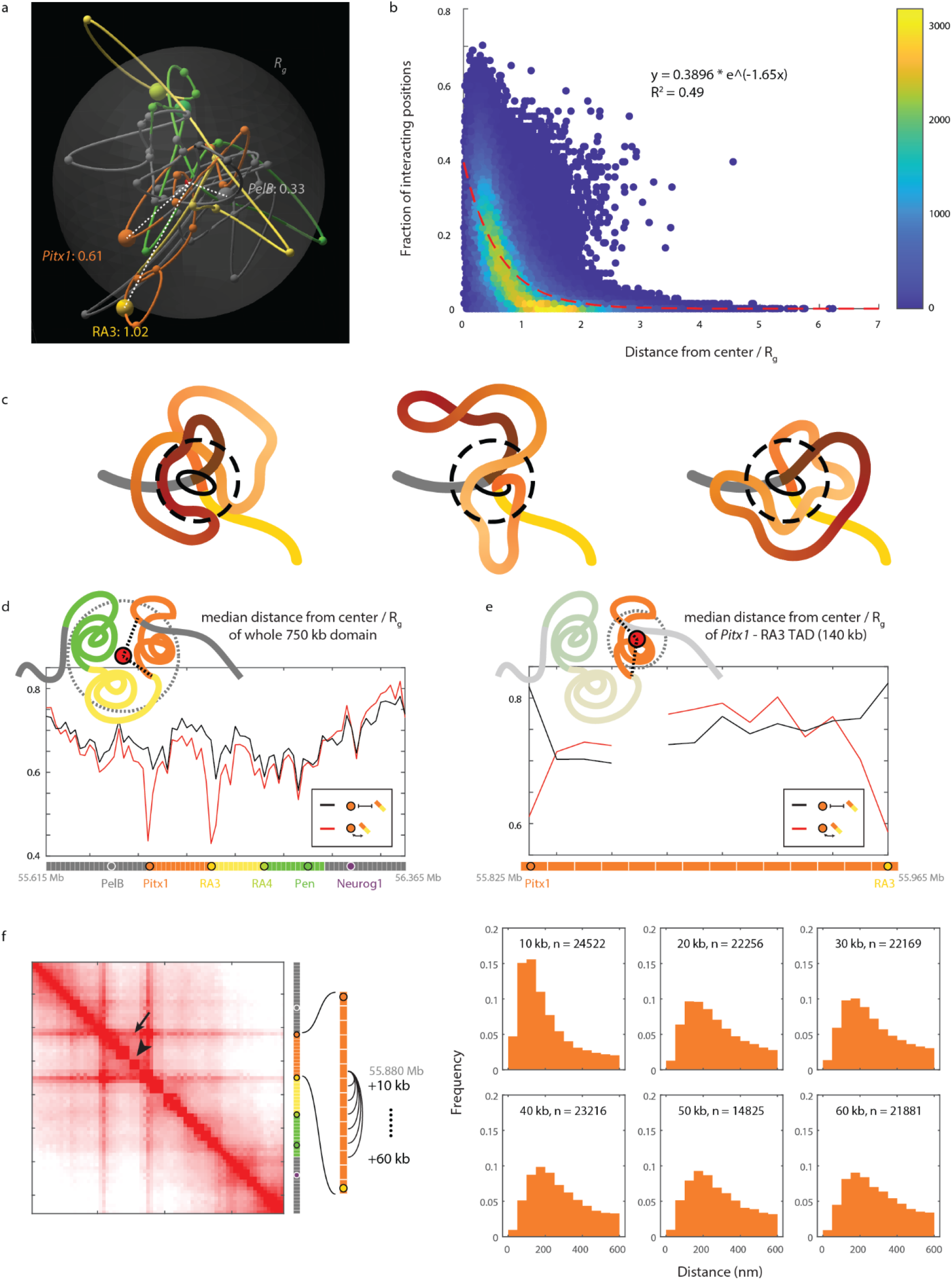
Central position within a molecule contributes significantly to contacting other parts of the molecule. **a,** Example chromatin trace showing how centrality is measured: for a position within the probed domain (colored spheres along the trace), we measure its distance from the center of the probed domain (small red sphere with radiating dotted white lines), normalized by the domain’s radius of gyration (*Rg*, translucent gray globe). The centrality of *PelB* (gray), *Pitx1* (orange) and RA3 (yellow) are shown as examples. **b,** The relationship between centrality within a molecule and contact frequency with other parts of the molecule, shown by data drawn from 641,687 10kb segments from 10,089 high-quality molecules across 5 independent experiments. Each dot represents a 10kb segment within the 750 kb probed domain. The X axis denotes distance from the center. The Y axis denotes the percentage of other segments within the same molecule contacting this segment (i.e. within 200 nm). Contacting multiple other segments happens mostly when a segment is positioned centrally in the molecule. Red dash line: exponential regression of the data, with its formula and R^2^ value indicated. **c,** Schematics that convey how central positioning can facilitate contact with all parts of the molecule. The dotted circles denote contact distance threshold. **d,** Distance from center of each 10kb segment across the *Pitx1* domain. Compared to all the molecules (black line), the subset of chromatin traces where the *Pitx1* promoter is contacting the RA3 enhancer (red line) are more likely to have both positions be at the center of the molecule. **e,** Even when considering the centrality only within the TAD between *Pitx1* and RA3, the two are still centrally positioned when they are in contact (red), despite their terminal linear positions. As illustrated by the cartoons on top of each subfigure, the differences between **d** and **e** are the choice of “center” (red) and radius of gyration (dashed gray circle). (Missing data: due to technical issues, the hybridization did not go well for the 5th data point in **e**. In other figures we interpolated for its position, but because doing so would result in a striking visual feature here, we left this segment blank). **f,** (Left) Contact frequency for molecules where *Pitx1* and RA3 are interacting, same as **Fig. 2c**. It might seem counterintuitive that being central to a region confers more contact than being linearly closer along the chromosome. (Compare the pixel marked by the arrow, representing interaction between two segments 80 kb apart, and that marked by the arrowhead, representing interaction between two segments 20 kb apart). However, what seems like a few pixel’s distance on the plot is actually many kilobases apart (since are regions tiled at 10 kb intervals), and within a TAD, which is not a rigid structure, the chromatin is rather flexible. (Right) Distributions of 3D distances between segment #27 (mm10, chr13: 55,875,000 - 55,885,000) and increasingly distant segments show that, despite being in the same TAD, distance distributions for segments over 30 kb apart do not differ much from each other, indicating that increased contact frequency conferred by linear proximity is limited to very proximal elements.

**Extended Data Fig. 5.**
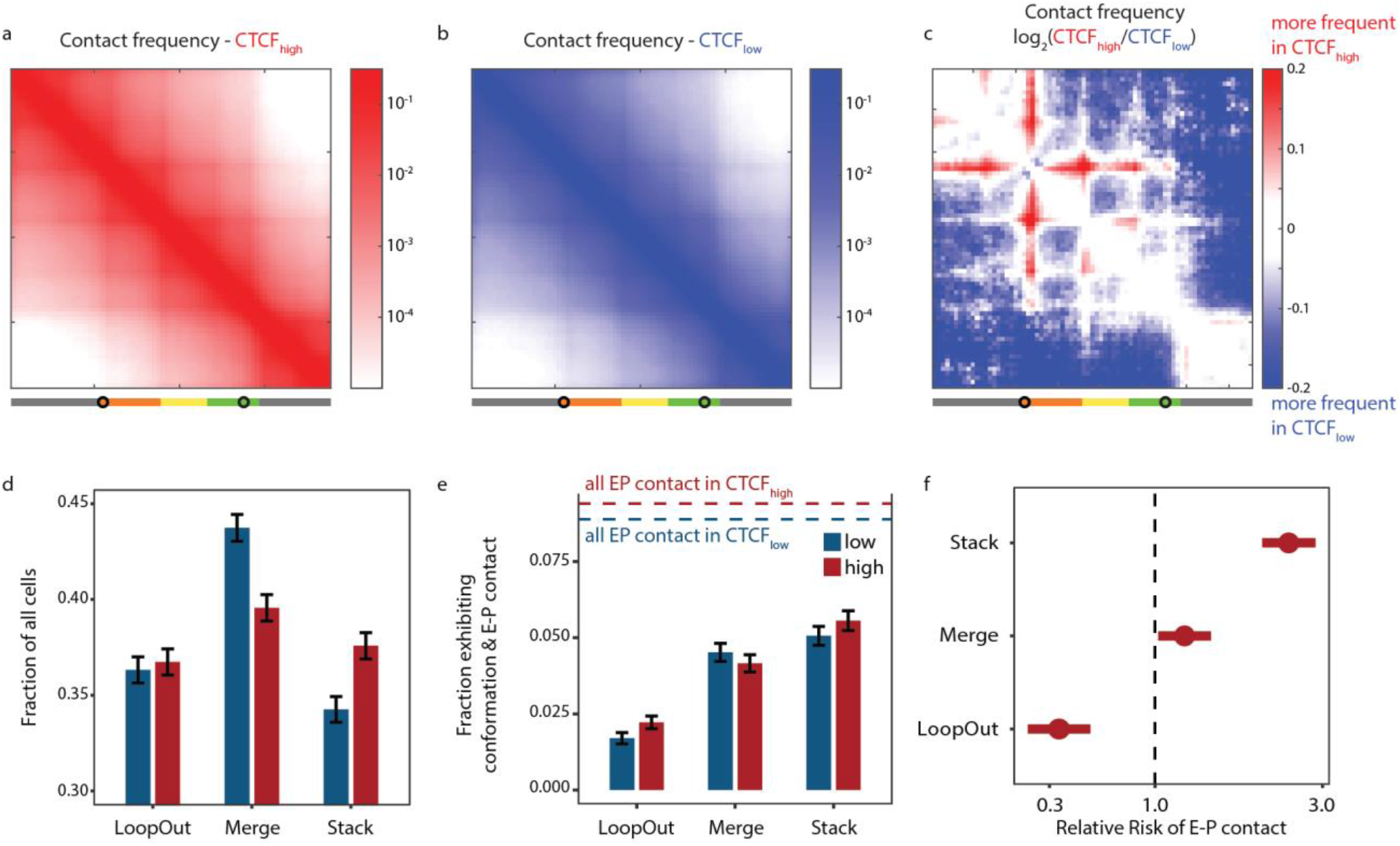
Modeling effects of increasing CTCF activity with fine-tuned parameters. **a,** Contact frequency at high CTCF. **b,** Contact frequency at low CTCF. **c,** log2 ratio of data in **a** and **b**. **d,** Fraction of all simulated polymers exhibiting configurations matching each model in low CTCF (blue) and high CTCF (red) conditions. Error bars represent standard deviations. Stack is the only configuration higher in the high CTCF condition. **e,** Fraction of all simulated polymers that have E-P interaction exhibiting configurations matching each model in low CTCF (blue) and high CTCF (red) conditions. Error bars represent standard deviations. Dotted lines represent the fraction of all polymers exhibiting E-P interaction. **f,** Relative risk of having E-P contact for polymers under high CTCF condition while exhibiting configurations matching a model. Bar indicates 95% confidence interval. Stack configuration grants the highest RR for EP contact. Although the RR for Merge is also positive, it is far more frequent in the low CTCF condition (see panel D).

## Supplementary Data

**Supplementary Table. 1.** Summary of the five experiments. Experiments 2 and 3 used different sections from the same limbs. Experiments 4 and 5 used different embryos from the same litter.

| ***Experiment#*** | **Stage** | **Left/Right** | **Barcodes probed** | **Forelimb N** | **Hindlimb N** |
| --- | --- | --- | --- | --- | --- |
| 1 | E12.5 | Right | 1-75 | 10087 | 11607 |
| 2* | E11.5 | Left | 1-75 | 7156 | 7326 |
| 3* | E11.5 | Left | 1-70 | 14220 | 3814 |
| 4 | E11.5 | Left | 1-75 | 8721 | 6459 |
| 5 | E11.5 | Right | 1-67 | 12992 | 15760 |

